# Prime editing in chicken fibroblasts and primordial germ cells

**DOI:** 10.1101/2022.05.31.494249

**Authors:** Yuji Atsuta, Katsuya Suzuki, Haruna Yaguchi, Daisuke Saito

**Affiliations:** Department of Biology, Faculty of Science, Kyushu University, Fukuoka 819-0395, Japan

**Keywords:** gene editing, prime editing, primordial germ cell, chicken

## Abstract

CRISPR/Cas9-based genome editing technologies are revolutionizing developmental biology. One of the advanced CRISPR-based techniques is prime editing (PE), which enables precise gene modification in multiple model organisms. In the current study, we describe a method to apply PE to the genome of chicken fibroblasts and primordial germ cells (PGCs). By combining PE with a transposon-mediated genomic integration, drug selection, and the single-cell culture method, we successfully generated prime-edited chick PGCs. The chicken PGC is widely used as an experimental model to study germ cell formation and as a vector for gene transfer to produce transgenic chickens. Such experimental models are useful in the developmental biology field and as potential bioreactors to produce pharmaceutical and nutritious proteins. Thus, the method presented here will provide not only a powerful tool to investigate gene function in germ cell development but also a basis for generating prime-edited transgenic birds.

## 1 INTRODUCTION

Recent advances in genome editing techniques based on the CRISPR/Cas9 system enable efficient and precise genomic modification of animal models. The latest representative, and perhaps most useful, technology is prime editing (PE), which enables genetic information to be inserted into a targeted genomic locus (Anzalone et al., 2019; Chen et al., 2021). The PE system consists of two operational components, a prime editor and a prime editing guide RNA (pegRNA). The prime editor is a fused protein of catalytically defective Cas9 endonuclease (dCas9 nickase) and engineered reverse transcriptase. The pegRNA is both an extended guide RNA containing spacer sequences to recognize and navigate dCas9 to a target site and a template for reverse transcription that encodes the desired edit. In comparison with previous CRISPR-based genome editing techniques, PE has several advantages, including avoidance of a double strand break, less constraint of the protospacer adjacent motif (PAM) location, and greater versatility than the base editing system (Komor *et al*., 2016; Rees & Liu, 2018; Scholefield & Harrison, 2021). Moreover, PE enables precise and efficient homology-directed repair in both mitotic and post-mitotic cells even without an additional repair donor construct (Anzalone *et al*., 2019).

Chicken primordial germ cells (PGCs) have been used as a model to study germ line development and cell migration (Stebler *et al*., 2004; Nakamura *et al*., 2007; Nakamura *et al*., 2013; Murai *et al*., 2021). In addition, because chicken PGCs can be expanded and gene-manipulated easily *in vitro* (Hong *et al*., 1998; Macdonald *et al*., 2012; Whyte *et al*., 2015), it has been used as a valuable vehicle of gene transfer to generate transgenic (TG) birds following the establishment of TG chickens by transplantation of genetically modified PGCs (van de Lavoir *et al*., 2006; Han & Park, 2018). TG chickens have great potential not only as a useful model for developmental biology but also as a bioreactor in the agricultural and pharmaceutical fields. Currently, CRISPR-based technologies, including base editing, have been used for genome modification of chicken PGCs (Dimitrov *et al*., 2016; Idoko-Akoh *et al*., 2018; Kim *et al*., 2020; Lee *et al*., 2020; Park *et al*., 2020). However, PE has not yet been applied to chicken PGCs.

Here, we tested PE for a transgene (*EBFP*) and endogenous gene (*DDX4*, a gene encoding a DEAD box RNA helicase) in chicken fibroblasts and PGCs. By taking advantage of a long-term culture system for chick PGCs and Tol2 mobile element-mediated genomic integration (Sato *et al*., 2007; Whyte *et al*., 2015), we devised a method to efficiently obtain clones of prime-edited PGCs. This method will open paths to investigating molecular mechanisms underlying PGC formation and germ cell development and to generating precisely gene-modified chickens.

## 2 MATERIALS AND METHODS

### 2.1 Experimental animals

Fertilized chicken eggs were obtained from Fujino-Kohkaen (Fukuoka, Japan). Embryos were staged according to the Hamburger–Hamilton stages (Hamburger & Hamilton, 1951). All animal experiments were conducted under the ethical approval of Kyushu University (No. A21-167-2).

### 2.2 Expression vectors and primers

EBFP-C1 (#54738), PX459 (#62988), BPK1520 (#65777), pU6-pegRNA-GG-acceptor (#132777), pCMV-PE2 (#132775), pCMV-PEmax (#174820), and pEF1a-hMLH1dn (#174824) were obtained from Addgene. pCAGGS-T2TP has been described elsewhere (Sato *et al*., 2007). For pT2A-CAGGS-EBFP, EBFP cDNA was amplified from EBFP-C1 by PCR with primers (EBFP-Fw, EBFP-Rv) and integrated into a MluI-NheI site of pT2A-CAGGS (Urasaki *et al*., 2006) using a Gibson Assembly Kit (New England BioLabs). To obtain pT2A-CAGGS-PEmax-ires-ZsGreen1, “PEmax” cDNA coding a fusion protein of optimized dCas9 and MMLV-reverse transcriptase was amplified from pCMV-PEmax using primers (PEmax-Fw, PEmax-Rv) and inserted into a MluI-EcoRI site of pT2A-CAGGS-ZsGreen1 (Atsuta & Takahashi, 2016) using a Gibson Assembly Kit. The pegRNAs used in this study were designed using pegFinder (http://pegfinder.sidichenlab.org; Chow *et al*., 2021). Template DNA (GFP pegRNA-1, GFP pegRNA-2, DDX4 pegRNA-1, DDX4 pegRNA-2, DDX4 pegRNA-3, DDX4 pegRNA-4, and DDX4 pegRNA-5) for pegRNA expression were synthesized and cloned into BsaI sites of pU6-pegRNA-GG-acceptor using a Gibson Assembly Kit. To obtain sgRNA-expressing PE3 plasmids (nicking vectors of the PE3 system), oligo nucleotides (PE3-EGFP-1-Fw and PE3-EGFP-1-Rv; PE3-EGFP-2-Fw and PE3-EGFP-2-Rv) were annealed and inserted into the BsmBI sites of BPK1520 using a Quick Ligation Kit (New England BioLabs). PuroR cDNA was amplified from PX459 with primers (PuroR-Fw, PuroP-Rv) and inserted into a MluI-EcoRI site of pT2A-CAGGS using a Gibson Assembly Kit, resulting in pT2A-CAGGS-PuroR. To construct the pT2A-U6-pegRNA-CAGGS-PuroR vector, U6-pegRNA and CAGGS-PuroR fragments were amplified with primers (U6-peg-Fw, U6-peg-Rv; CAG-PuroR-Fw, CAG-PuroR-Rv) from pegRNA vectors and pT2A-CAGGS-PuroR, respectively. The fragments were cloned into the ApaI-MluI and MluI-BglII sites of pT2A vectors using Ligation High ver.2 (TOYOBO), resulting in the construction of pT2A-U6-pegRNA-CAGGS-PuroR plasmids. Sequences of the primers, template DNA for pegRNA expression, and oligos for sgRNA-expressing PE3 plasmids are shown in Supplemental Table 1.

### 2.3 Cell culture and derivation of PGCs

Chicken fibroblast-derived DF1 cells were maintained at 38°C with DMEM/F12 (Gibco) containing 10% fetal bovine serum (Biowest) and 1% penicillin–streptomycin (Gibco). PGCs were obtained and maintained as previously described (Whyte *et al*., 2015), with minor modifications (Chen *et al*., 2019). PGCs were derived by transferring 1–2 μL blood isolated from HH16 embryos in 300 μL of media (FAcs media) composed of DMEM (Ca^2+^ free, Gibco), 100 μM CaCl_2_, 1× B-27 supplement (Gibco), 2 Mm Glutamax (Gibco), 1× NEAA (Gibco), 55 μM β-mercaptoethanol (Gibco), 1.2 mM pyruvate (Gibco), 0.2% chick serum (Biowest), 0.2% ovalbumin (Sigma-Aldrich), 0.2% heparin (Sigma-Aldrich), 25 ng/mL of human activin A, and 4 ng/mL human FGF2 (R&D). PGCs were basically maintained in the FAcs medium at 38°C.

### 2.4 DNA transfection, FAC-sorting and PGC electroporation

DF1 cells were seeded in a 48 well plate and transfected at 60–70% confluency with Lipofectamine 2000 (Thermo Fisher Scientific). To establish EBFP-expressing DF1 cells, 1 μL Lipofectamine 2000, 750 ng pT2A-CAGGS-EBFP, and 250 ng pCAGGS-T2TP were transfected. One week after transfection, cells were dissociated with 100 μL 0.05% trypsin–EDTA solution (Gibco), and EBFP-expressing cells were collected using an SH800 cell sorter (Sony). For PE in DF1 cells, 1 μL Lipofectamine 2000, 500 ng prime editor plasmid, 150 ng pegRNA plasmid, 50 ng sgRNA plasmid (a nicking plasmid for PE3; where indicated), and 150 ng pEF1a-hMLH1dn (where indicated) were used. The efficiency of EBFP-to-EGFP conversion was estimated using an SH800 cell sorter. To obtain the PEmax expressing DF1 cells, 1 μL Lipofectamine 2000, 750 ng pT2A-CAGGS-PEmax-ires-ZsGreen1, and 250 ng pCAGGS-T2TP were transfected and ZsGreen-positive cells were sorted using an SH800 cell sorter 1 week after transfection. For the second transfection to produce *DDX4* edited cells, pT2A-U6-pegRNA-CAGGS-PuroR and pEF1a-hMLH1dn were further introduced into PEmax-expressing cells. The cells were maintained in the presence of 1 μg/mL puromycin (Nacalai Tesque) for 3 days, then washed to remove puromycin and further cultured to 100% confluency. For electroporation to PGCs, 2 × 10^4^ PGCs were resuspended in 10 μL R buffer containing 1 μg DNA plasmid (750 ng pT2A-CAGGS-EBFP, 250 ng pCAGGS-T2TP; 600 ng pCMV-PEmax, 300 ng GFP pegRNA-1 plasmid, 100 ng PE3-EGFP-1 plasmid; 750 ng pT2A-CAGGS-PEmax-ires-ZsGreen1, 250 ng pCAGGS-T2TP; 450 ng DDX4 pegRNA-4 plasmid, 100 ng PE3-DDX plasmid, 450 ng pEF1a-hMLH1dn) and electroporated using a Neon transfection system (Thermo Fisher Scientific) with three pulses of 1300 V for 10 ms. To select pegRNA-transfected cells, PGCs were selected using puromycin for 3 days, followed by 1 week of puromycin-free culture. For a clonal culture, individual cells were sorted in a 96 well plate using an SH800 cell sorter after puromycin selection and were then further cultured for 28 days. Genomic DNA extraction for the prime edited DF1 or PGCs was carried out with NucleoSpin Tissue XS (TAKARA) according to the manufacturer’s protocol. To confirm the genomic edit, a portion of the *DDX4* locus (chrZ:16,929,487-16,929,972) was amplified by PCR with a set of primers (DDX4-seq-Fw, DDX4-seq-Rv) and the PCR products were sequenced using the DDX4-seq-Fw primer.

### 2.5 Embryo injection of PGCs

EGFP-expressing PGCs (3000–5000 cells/μL) were injected into hearts of HH14 host embryos using a glass capillary and incubated until HH26 or HH30. Gonads of the host embryos were subject to immunological staining.

### 2.6 Immunohistochemistry

For immunological staining on sections, the following primary antibodies were used: chick anti-GFP (1/1000; abcam, ab13970), rat anti-DDX4 (1/1000; (Yoshino *et al*., 2016)), and rabbit anti-Sox9 (1/500, Millipore, AB5535). Immunostaining for chicken embryos was carried out as previously described (Tomizawa *et al*., 2021). The embryos were fixed in 4% PFA/PBS for 3 h at 4°C, placed in a series of sucrose/PBS solutions, and embedded in OCT compound (Sakura Finetek). Ten micrometer cryo-sections were permeabilized with 0.5% Triton X-100/PBS for 10 min and then incubated with blocking buffer (1% blocking reagent [Roche]/TNT) for 1 h, followed by incubation of primary antibodies overnight at 4°C. Sections were washed with TNT buffer and incubated with secondary antibodies for 5 h at 4°C. Alexa 488 anti-chicken IgY (abcam), Alexa 568 anti-rat IgG, Alexa 568 anti-rabbit IgG, and Alexa 488 anti-mouse IgG (Thermo Fisher Scientific) were used as secondary antibodies. Vectashield mounting media with DAPI (Vector Laboratories) was used before imaging. The sex of HH30 gonads were determined by Sox9 positivity. For immunostaining with cultured PGCs, the cells were incubated with anti-DDX4 antibody after PFA fixation and permeabilization of 0.5% Triton X-100/PBS, followed by incubation of Alexa 568 anti-rat IgG.

### 2.7 Imaging and statistical analysis

Images of stained sections were obtained using a Dragonfly confocal microscope (Andor). Statistical analyses were performed with GraphPad Prism 9 software (GraphPad). The live cells in Fig. 4D were counted by using an automated cell counter (TC20, Bio-Rad), Counting slides (Bio-Rad), and trypan blue (FUJIFILM).

## 3 RESULTS AND DISCUSSION

### 3.1 PE is applicable to the chicken cell line

To test whether PE can be used in avian cells and to efficiently visualize successful genomic modification by PE, we set up the EBFP–EGFP conversion system with the DF1 chicken fibroblast cell line. There are two nucleotide substitutions (T199C, A437T) that result in amino acid substitutions (Y66H, Y145F) between DNA sequences encoding enhanced green fluorescent protein (*EGFP*) and the enhanced blue fluorescent protein (*EBFP*) (Heim & Tsien, 1996) (Fig. 1A). One of the substitutions, designated as Target site-1 in the present study, is responsible for the determination of the fluorescent spectrum, whereas the other, Target site-2, enhances the brightness of blue fluorescence (Heim *et al*., 1994; Heim & Tsien, 1996) (Fig. 1A). Thus, we expected that, if Target site-1 C^199^ was replaced with thymidine using the PE technique, a successful replacement could be monitored via EGFP fluorescence. We generated the stable EBFP DF1 cell line by taking advantage of Tol2 transposon-mediated genomic integration (Urasaki *et al*., 2006; Sato *et al*., 2007) (Fig. 1B). Prime editor- and pegRNA-1 (pegRNA aiming Target site-1)-expressing vectors were transfected to the EBFP cells. As a result, EGFP-expressing cells emerged 2 days after transfection, indicating that the PE technique is functional in chicken cells, as observed in other organisms (PE2; Fig. 1B). Editing Target site-2 generated no EGFP-positive cells, consistent with the results of a previous study (Fig. 1C) (Heim & Tsien, 1996).

**FIGURE 1.**
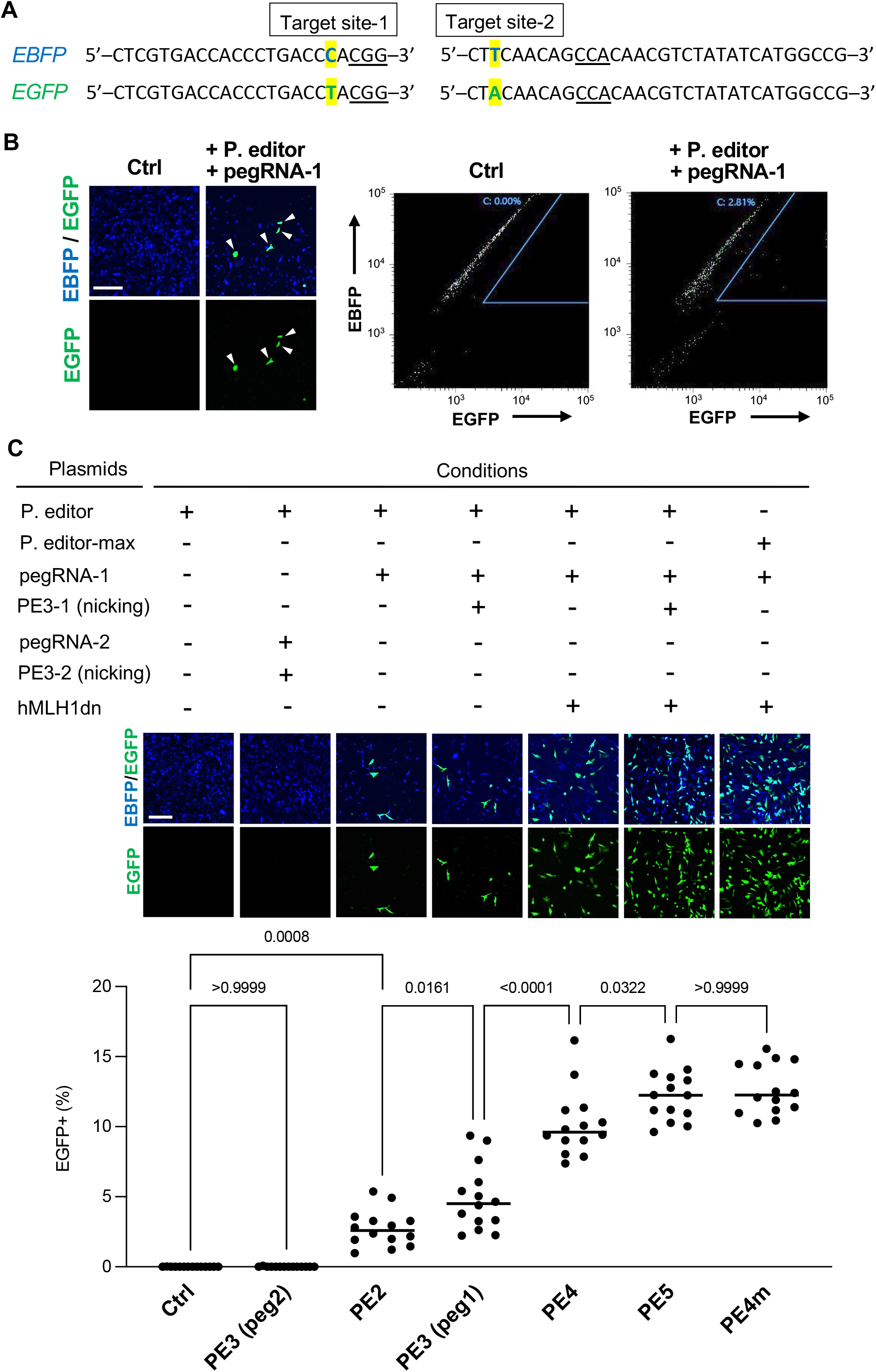
Prime editing for transgenes in chicken fibroblast cells. (A) Target sites for EBFP–EGFP conversion by prime editing (PE). PAM sequences are underlined. (B) EBFP-expressing DF1 cells were transfected with prime editor- and pegRNA-1 (targeting site-1)-expressing plasmids (PE2 plasmids), and a small fraction of the cells started showing EGFP fluorescence (arrowheads). The fluorescence was validated by FACS. (C) Comparison of different PE conditions for the EBFP/EGFP conversion in DF1 cells (*n* = 14 wells for each condition). PE3 vectors are nicking plasmids. The numbers show *P* values obtained using ordinary one-way ANOVA. Scale bars: 200 μm in (B, C).

Improved versions of PE that enhanced editing efficiency by manipulating the DNA repair machinery have recently been reported (Chen *et al*., 2021). Therefore, we used the EBFP–EGFP conversion system to examine which version of the PE system was most effective in chicken cells (Fig. 1C). As expected, the EBFP–EGFP conversion efficiency of the PE3 system (PE2 plus a nicking vector) was higher than that of the PE2 system (the prime editor vector plus a pegRNA expressing vector) (Fig. 1C). We further tested two engineered versions of PE: PE4 and PE5 (Chen *et al*., 2021). PE4 is composed of PE2 and a dominant negative form of DNA mismatch repair protein (MLH1dn)-expressing vector, whereas PE5 is a combined system of PE3 and MLH1dn. As shown in Fig. 1C, the addition of MLH1dn substantially increased the conversion efficiency of PE2 and PE3 (Fig. 1C), suggesting that DNA repair machinery diminishes PE efficacy in chicken cells, consistent with its effect in other animal cells (Chen *et al*., 2021). We also used a PEmax vector expressing an optimized prime editor architecture, instead of the basic one, in the PE4 system (referred to as PE4m) (Fig. 1C). The PE4m system yielded comparable efficiency as the PE5 system even in the absence of a nicking vector (Fig. 1C). Thus, the PE4m system was used in subsequent experiments.

### 3.2 PE does not the affect migratory behavior of PGCs

One of the major advantages of PE is that it leads to fewer off-target effects compared with conventional CRISPR/Cas9, thereby rarely causing phenotypic alteration (Anzalone *et al*., 2019). Because we aimed at applying PE to chicken PGCs, we investigated whether PE affected PGC character by checking the migratory capability of prime-edited PGCs (Fig. 2). EGFP-positive PGCs generated using the Tol2 system have been shown to migrate correctly toward the genital ridge when injected into the circulation of host embryos (Macdonald *et al*., 2012). EGFP-expressing PGCs derived from EBFP-positive cells were sorted and propagated in an optimal medium (FAcs medium, see MATERIALS AND METHODS), then injected into HH14 host embryos (Fig. 2A, B). We found that the grafted PGCs could reach the genital ridge of HH26 embryos and the developing gonads of both male and female HH30 embryos (Fig. 2B). These results suggest that PE does not affect the migratory behavior at least, which is one of the most representative traits of chicken PGCs.

**FIGURE 2.**
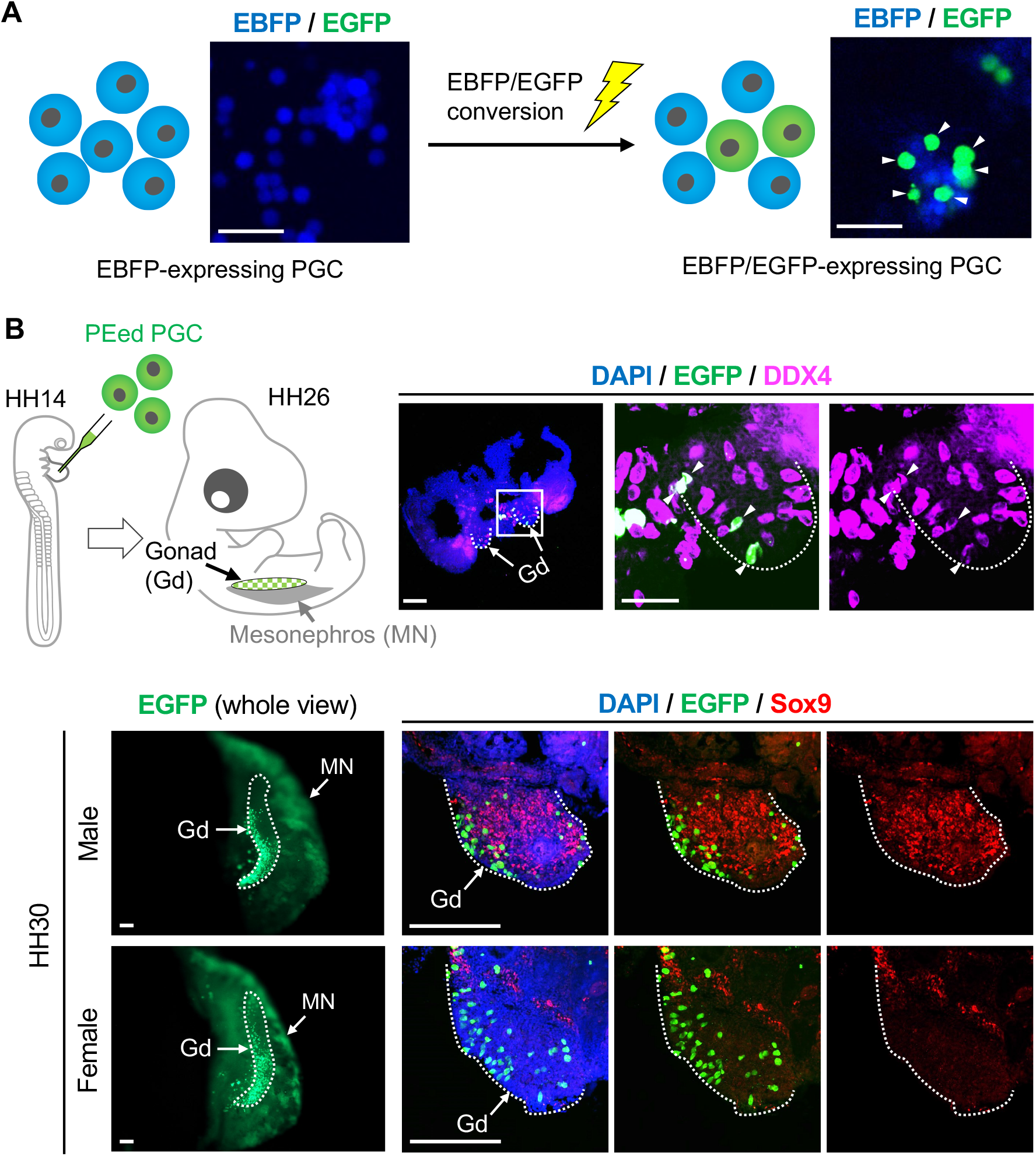
Grafted prime-edited PGCs migrate correctly to the genital ridge in chicken embryos. (A) *EBFP* genes in EBFP-expressing PGCs were converted to *EGFP* by prime editing (PE4m). (B, C) The PEed EGFP-expressing PGCs were sorted and subsequently transplanted into HH14 embryos. The grafted cells were found at the genital ridges of HH26 (B; *n* = 4 embryos) and HH30 male and female embryos (C; *n* = 3 for each sex). Scale bars: 50 μm in (A), 100 μm in (B), 200 μm in (C).

### 3.3 Base substitution of *DDX4* gene by PE in the chicken cell line

Having verified that transgenes in the chicken genome can be modified by PE, we next attempted to edit DNA sequences for endogenous genes. The *vasa* gene, which was originally isolated in *Drosophila* (Schupbach & Wieschaus, 1986), encodes a DEAD box RNA helicase, and chicken *vasa* homologue *cDDX4* (also referred to as *CVH*) is a prominent marker gene for PGCs (Tsunekawa *et al*., 2000). Thus, we speculated that *DDX4* could be a candidate to validate PE for an endogenous gene in chicken PGCs. We first experimented with DF1 cells because of their high availability (Fig. 3). To elevate editing efficiency and to concentrate gene-edited cells, we derived DF1 cells that stably expressed PEmax and ZsGreen1 bicistronically (Fig. 3A). We also used a puromycin selection to harvest only pegRNA-expressing cells (Fig. 3A). Considering the sequence preferences of guide RNAs in the CRISPR/Cas9 system (Xu *et al*., 2015), we designed multiple pegRNAs: pegRNA-1, -2, and -3 to abrogate the start codon of *DDX4* and pegRNA-4 and -5 to install a stop codon downstream of the start codon (Fig. 3B). Genomic DNAs were extracted from DF1 cells transfected with PE components, and the targeted regions were sequenced (Fig. 3B). We observed peaks that originated from mutated alleles of the transfected cells, except in the case of pegRNA-5-transfected cells (Fig. 3B). Among pegRNA-1 to -4, pegRNA-4 appeared to be efficient, and the substitution (G → T) by pegRNA-4 could not be installed by base editing techniques already applied to genome modification in chickens (Komor *et al*., 2016; Lee *et al*., 2020; Porto *et al*., 2020). Therefore, pegRNA-4 was chosen to edit DDX4 in PGC in subsequent experiments.

**FIGURE 3.**
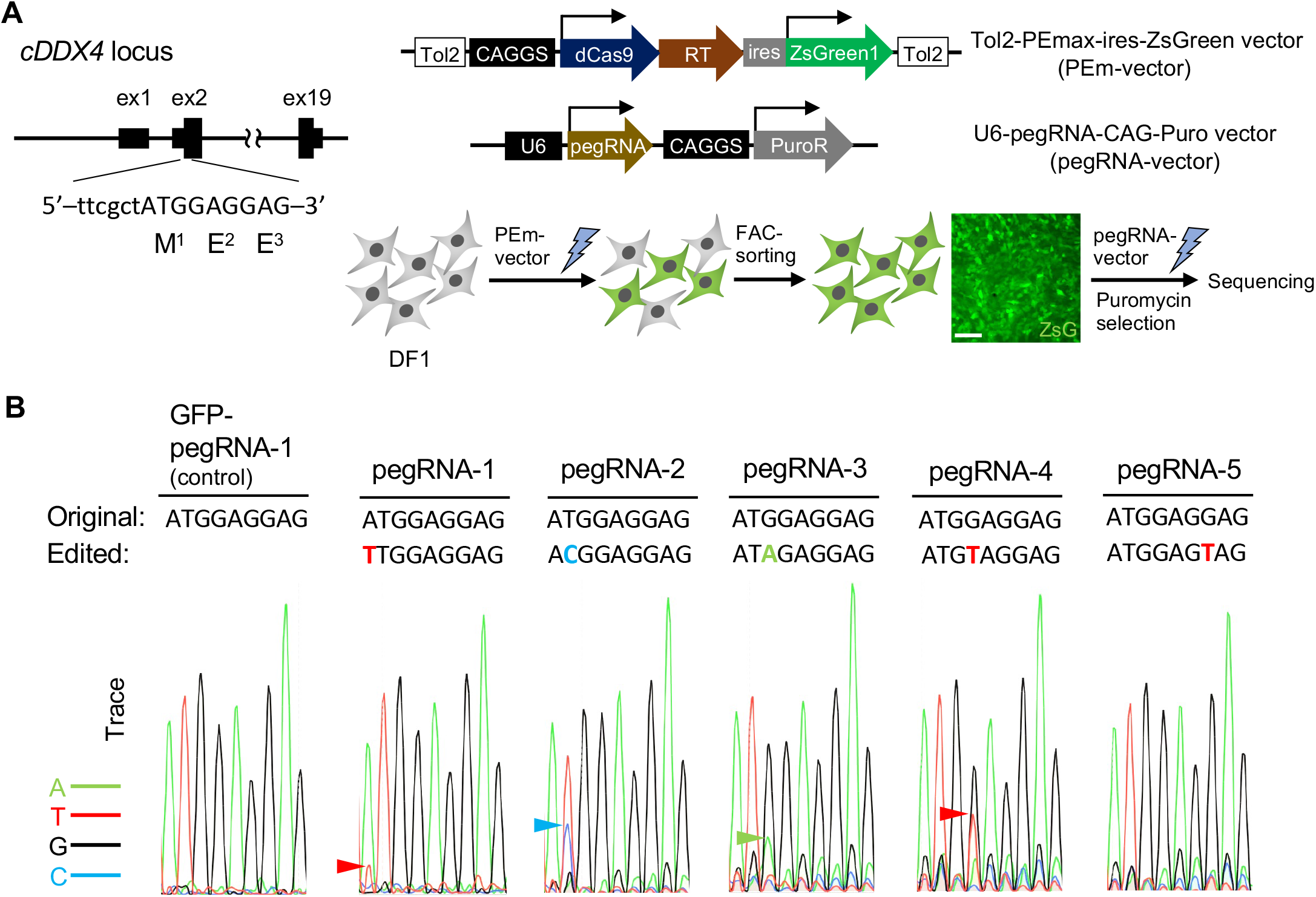
Prime editing for an endogenous gene in chicken fibroblast cells. (A) Schematics for *DDX4* locus and strategy for editing *DDX4* in DF1 cells. The *DDX* gene lies at chrZ: 16,929,379–16,960,987. Cells that stably express PEmax and ZsGreen1 (ZsG) were established by taking advantage of Tol2-mediated genomic integration, and the ZsG^+^ cells were FAC-sorted. PegRNA-transfected cells were further selected by puromycin. After the selection, DNAs extracted from transfected cells were subject to Sanger sequencing. (B) Chromatograms of DNA sequences coding for the first three codons of *DDX4*. PegRNAs were designed to abrogate the first codon ATG (pegRNA-1, -2, and -3) or to generate a stop codon after ATG (pegRNA-4 and -5). Peaks for the edited nucleotides are indicated by arrowheads. Scale bar: 100 μm in (A).

**FIGURE 4.**
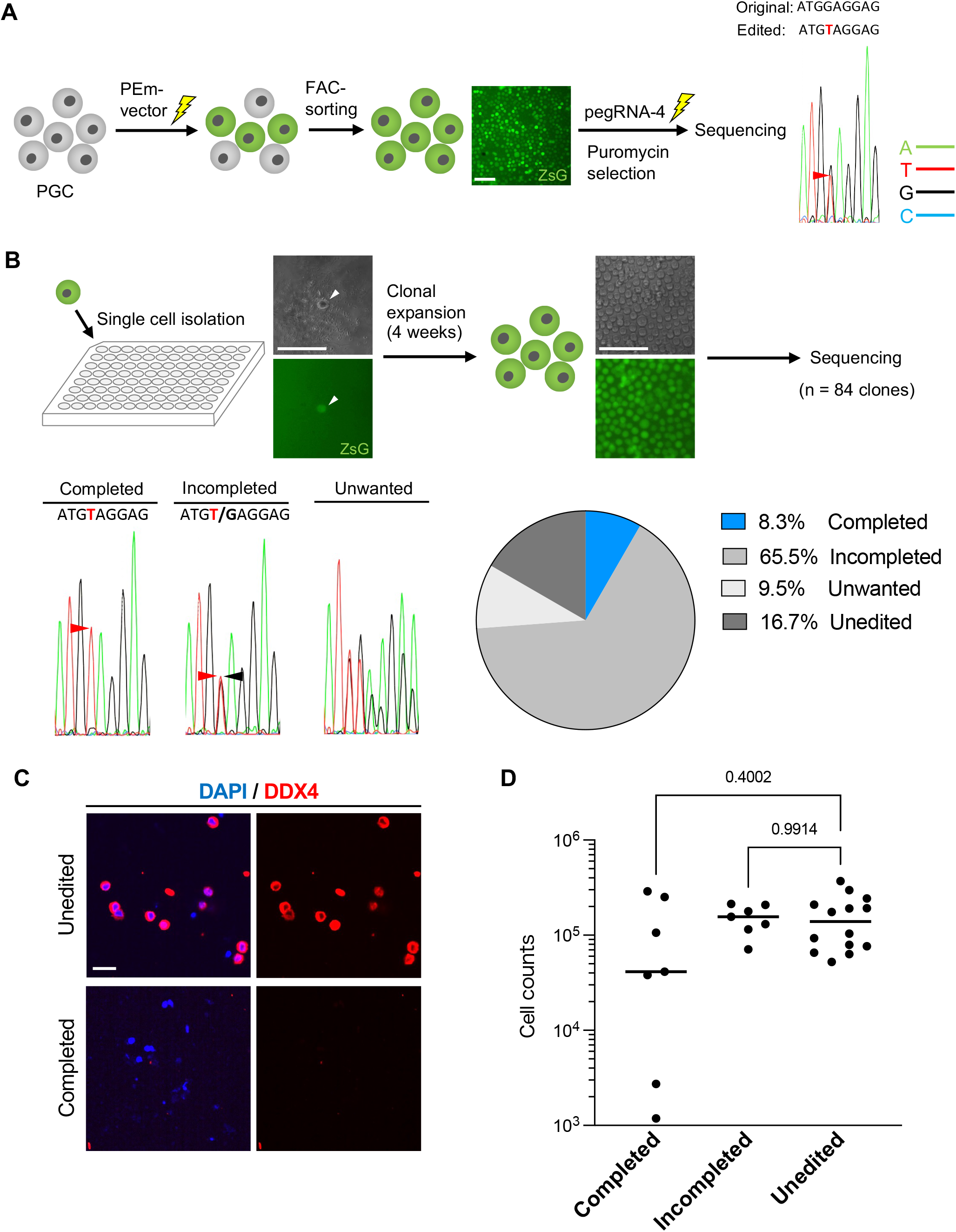
Isolation of the prime-edited PGC clones. (A) PEmax and ZsGreen-expressing PGCs were generated and collected using a cell sorter. After transfection of pegRNA-4 vector and puromycin selection, DNAs were extracted from the transfected PGCs and sequenced to validate the successful edit (red arrowhead). (B) Schematic of experiments for identifying the edited PGC clones. Individual cells were isolated using a sorter and cultured for 28 days before sequencing. Traces represent completely, incompletely, and unwantedly edited DNA sequences (“Completed,” “Incompleted,” and “Unwanted,” respectively). The pie chart shows the percentage of each genotypic group. (C) “Unedited” and “Completed” PGCs were immunostained against DDX4. DDX4 proteins were not detected in mutated cells. (D) Cell numbers were measured 28 days after the single-cell isolation (*n* = 7 wells for “completed,” *n* = 7 wells for “incompleted,” *n* = 14 wells for “unedited”). The numbers indicate *P* values obtained using an ordinary one-way ANOVA. Scale bars: 100 μm in (A, B), 20 μm in (C).

### 3.4 Development of a method to isolate prime-edited PGC clones

Lastly, to isolate desired mutant PGC clones, we established a combined method of PE4m and a single-cell culture system (Fig. 4). PGCs transfected with both PEmax and pegRNA-4 vectors were obtained by taking advantage of FAC-sorting and puromycin selection, as performed with DF1 (Fig. 4A). After we confirmed the substitution, we isolated the PGCs individually and expanded them in the FAcs medium for 4 weeks until sequencing analyses (Fig. 4B). Among 84 clones obtained, 7 clones had the desired substitution (8.3%), whereas the substitution was not detected in 14 clones (16.7%; Fig. 4B). We observed nucleotide substitutions in the majority of the clones; however, most of them were incomplete (65.5%) or undesired (16.7%) edits (Fig. 4B). Nevertheless, our method successfully identified and isolated the PGC clones that harbored the precise, desired substitution.

We further examined whether *DDX4* was indeed knocked out by inserting a stop codon into the locus and whether the stop codon affected the proliferation of PGCs. By using the antibody that specifically recognizes the N-terminus of *DDX4* (Raucci *et al*., 2015, Yoshino *et al*., 2016), we found that *DDX4* proteins were not translated in mutant cells, indicating the successful disruption of *DDX4* (Fig. 4C). To assess whether *DDX4* played a pivotal role in PGC proliferation, we counted the number of *DDX4*-KO cells. No significant difference was observed in the cell numbers between the unedited and edited groups (Fig. 4D), implying that *DDX4* is dispensable for mitosis of chicken PGC even though it is essential for meiotic processes during germ cell specification in multiple species (Kuramochi-Miyagawa *et al*., 2010; Ewen-Campen *et al*., 2013; Hartung *et al*., 2014; Taylor *et al*., 2017).

We successfully applied the PE technique to the chicken fibroblast cell line and PGCs. Combined with Tol2 and the single-cell culture systems, this application enabled us to obtain the desired mutant PGC clones. The proposed method would provide precise genome editing in PGCs and enable tests for gene function in PGC development in both *in vitro* and *in vivo* settings. Furthermore, because the PGC-mediated germline transmission system is widely used to generate transgenic chickens (van de Lavoir *et al*., 2006; Macdonald *et al*., 2010; Macdonald *et al*., 2012; Oishi *et al*., 2016; Kim *et al*., 2020; Lee *et al*., 2020; Park *et al*., 2020), our method could underlie potential applications for the efficient genomic modification of chickens.

## ACKNOWLEDGEMENTS

We thank Dr. Ryan Delgado (Harvard Medical School) for helpful discussions and the Center for Advanced Instrumental and Educational Support of the Faculty of Agriculture (Kyushu University) for the use of the SH800. This work was supported by the JST FOREST Program, Grant Number JPMJFR214G (to Y.A.), and JSPS KAKENHI Grant Number JP20K22658, JP21K06201 (to Y.A.), and JP18H02445 (to D.S.).

## AUTHOR CONTRIBUTIONS

**Yuji Atsuta:** study design; formal analysis; investigation; methodology; writing-original draft; writing-review & editing; funding acquisition. **Katsuta Suzuki:** investigation; methodology; writing-review & editing. **Haruna Yaguchi:** investigation; methodology; writing-review & editing. **Daisuke Saito:** investigation; methodology; writing-review & editing; funding acquisition. All authors reviewed and approved the manuscript.

---

**Table S1.**
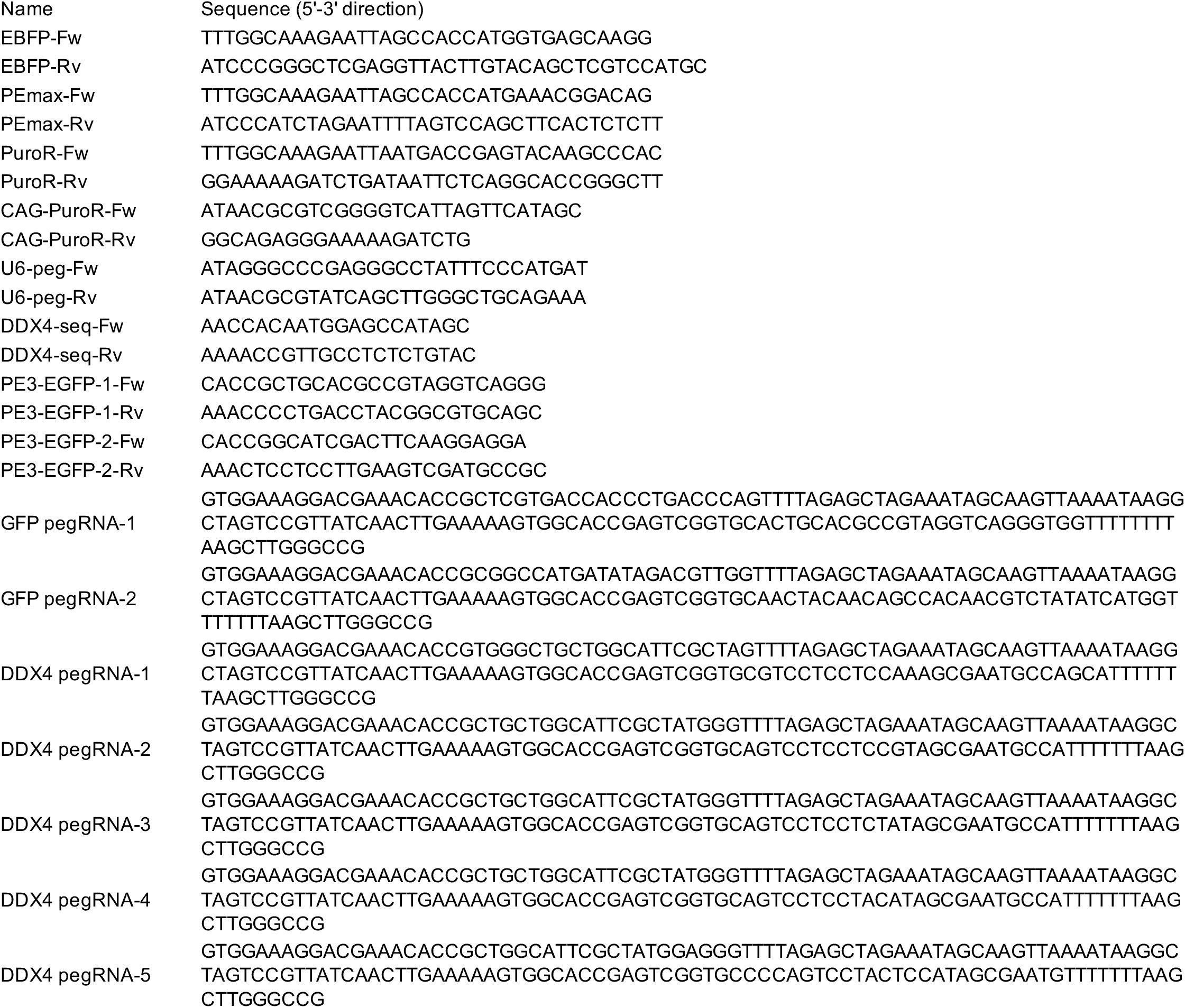
Sequences of primers and pegRNA templates.

